# *FBXO42* activity is required to prevent mitotic arrest, spindle assembly checkpoint activation, and lethality in glioblastoma and other cancers

**DOI:** 10.1101/2022.11.29.518420

**Authors:** Pia Hoellerbauer, Megan Kufeld, Sonali Arora, Emily J. Girard, Jacob A. Herman, James M. Olson, Patrick J. Paddison

## Abstract

Glioblastoma (GBM) is the most common and aggressive brain tumor in adults. To identify genes differentially required for the viability of GBM stem-like cells (GSCs), we performed functional genomic lethality screens comparing GSCs and control human neural stem cells. Among top scoring hits in a subset of GBM cells was the F-box-containing gene *FBXO42*, which was also essential in ∼15% of cell lines derived from a broad range of cancers. Mechanistic studies revealed that, in sensitive cells, *FBXO42* activity prevents chromosome alignment defects, mitotic cell cycle arrest, and cell death. The cell cycle arrest, but not the cell death, triggered by *FBXO42* inactivation could be suppressed by brief exposure to a chemical inhibitor of Mps1, a key spindle assembly checkpoint (SAC) kinase. *FBXO42*’s cancer-essential function requires its F-box and Kelch domains, which are necessary for FBXO42’s substrate recognition and targeting by SCF ubiquitin ligase complex. However, none of FBXO42’s previously proposed targets, including ING4, p53, and RBPJ, were responsible for the observed phenotypes. Instead, our results suggest that *FBOX42* activity suppresses the accumulation of one or more proteins that perturb chromosome-microtubule dynamics in cancer cells, which, in turn, leads to induction of the SAC and cell death.

## Introduction

Gliomas account for approximately 30% of all central nervous system tumors [1]. The most aggressive and common form is glioblastoma (GBM)[2]. There are currently no highly effective therapies against GBM. With standard-of-care (SOC) treatments, consisting of surgery, radiation, and the alkylating agent temozolomide, ∼90% of adult patients die within 2 years of diagnosis [3, 4]. The median survival for GBM patients overall ranges from 14-17 months, with rare exceptions of long-term survival [5, 6].

Although SOC effectively “debulks” GBM tumors, it doesn’t prevent tumor regrowth and disease recurrence, resulting in the observed modest survival benefits. The prevailing rationale is that tumors harbor slow dividing GBM stem-like cells (GSCs), which are both missed by surgery and SOC resistant and cause tumor regrowth [7-10]. This concept was elegantly demonstrated in a mouse model of glioma where a quiescent subset of endogenous glioma cells was shown to be responsible for tumor regrowth after TMZ treatment [11].

For human GBM, serum-free culture methods that resemble a stem cell niche allow retention of many of the properties associated with patient tumor isolate, including stem-like cell states [7-10]. Using these serum-free culture conditions, we have previously performed functional genomic screens in adult GBM-stem-like cells (GSCs), comparing them to human neural stem cells (NSCs) to identify cancer-specific hits [12-17]. These screens identified multiple cancer-specific dependencies, including: the GLEBS domain of BubR1/BUB1B, which suppresses Ras/MAPK-driven kinetochore-microtubule attachment defects [12, 13, 18]; *PHF5A*, which helps maintain proper exon recognition in MYC overexpressing cells [14]; *BuGZ*, which facilitates Bub3 activity and chromosome congression [15]; *PKMYT1*, which prevents premature entry into mitosis in response to over activity RTK/PI3K signaling [16]; and *ZNF131*, which promotes expression of HAUS5 to maintain the integrity of centrosome function in GBM cells [17].

Here, we characterize another hit from CRISPR-Cas9 lethality screens performed in GSCs, *FBXO42*, which shows cancer-specific requirement to prevent mitotic arrest, spindle assembly checkpoint activation, loss of viability, and loss of tumor maintenance.

## Results

### CRISPR-Cas9 screen identify *FBXO42* as a selective lethal gene in a subset of GSC isolates

We previously performed genome-scale CRISPR-Cas9 lethality screens in two adult human GSCs (GSC-0131-mesenchymal and GSC-0827-proneural) and two human neural stem cell (NSC) isolates (CB660 and U5) to identify genes differentially required for GSC outgrowth isolates [16] (Figure 1A). Among the strongest GSC selective hits screens was *FBXO42*, which scored as essential in GSC-0827 cells, but was seemingly completely dispensable in GSC-0131, GSC-1502, and NSCs. *FBXO42* encodes an F-box protein that serves as the substrate-recognition component of an SCF (SKP1-CUL1-F-box protein)-type E3 ubiquitin ligase complex, and it is thus far poorly characterized, with only a few publications exploring the roles of the protein [19-22]. One previous study described a role for FBXO42 in the destabilization of p53, similar to MDM2 [20]. Because GSC-0827 cells are *TP53*-wt and GSC-0131 are *TP53*-mutant, we wondered if *TP53* status might explain *FBXO42*’s GSC-0827-specific requirement. We, therefore, decided to explore this screen hit further.

**Figure 1:**
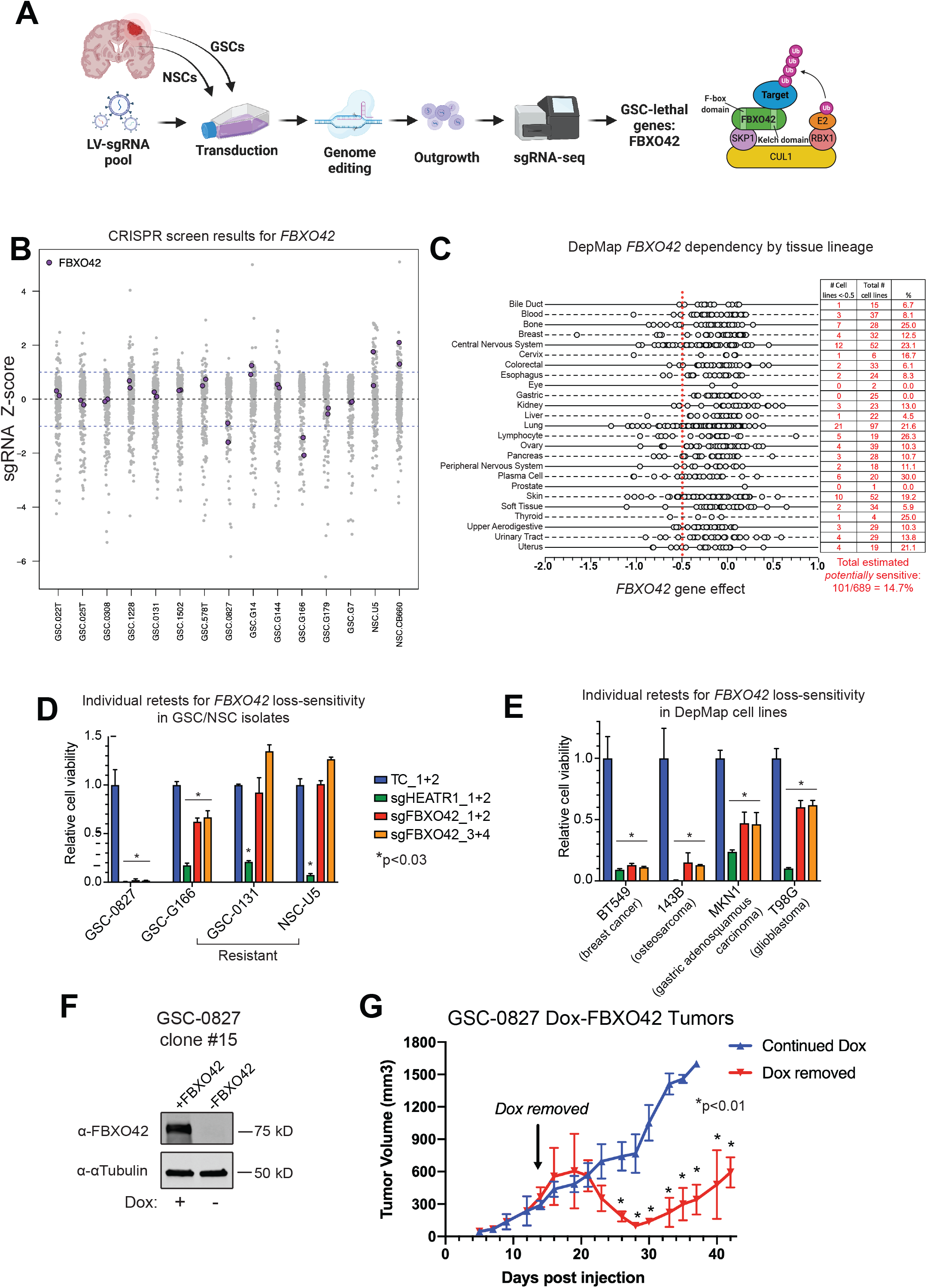
Identification of *FBXO42* as a candidate cancer-lethal gene. **A**, Overview of functional genomic screens that revealed *FBXO42* as a candidate GBM-lethal gene. **B**, Retest screens of GSC-specific hits from Toledo et al., 2015, showing how sgRNAs targeting *FBXO42* scored. Cells were infected with LV-Cas9-sgRNA pool, outgrown for 21 days, and subjected to sgRNA-seq. **C**, Breakdown by tissue lineage of *FBXO42* dependency in cell lines screened in DepMap (CERES score from CRISPR Avana 19Q4 dataset). Each circle corresponds to a cell line. Summary of proportion of potentially sensitive cell lines (score <-0.5) by tissue lineage is shown to the right. **D**, Relative cell viability (normalized to targeting control *sgCD8A*) for GSCs and NSCs nucleofected with CRISPR RNPs targeting *FBXO42. HEATR1* is an essential control gene. *sgNTC* = non-targeting control sgRNA. Measured at 9 days post nucleofection for GSCs and 11 days for NSCs (due to doubling time differences). **E**, Relative cell viability (normalized to targeting control *sgCD8A*) for 4 cell lines that were nucleofected with CRISPR RNPs targeting *FBXO42*. Lines had been predicted *FBXO42* loss-sensitive based on DepMap data (using CRISPR Avana dataset for all except MKN1, which were predicted based on Combined RNAi dataset.) *HEATR1* is an essential control gene. *sgNTC* = non-targeting control sgRNA. Measured at 8-12 days post nucleofection (depending on doubling time). **F**, Western blot for FBXO42 for clone 15 with either continuous doxycycline exposure, 4 days doxycycline removal. Clone 15 cells harbor a biallelic disruption of endogenous *FBXO42* locus and also express a Dox-controlled *FBXO42* ORF, which complements the loss of endogenous *FBXO42*. **G**, Tumor growth of GSC-0827 Dox-*FBXO42* clone 15 cells with and without doxycycline. Flank tumors (n= 5 for each arm; *pval <.01, student’s t-test) were used to avoid any potential bioavailability issues with dox crossing the blood brain barrier.

First, we performed a retest screen among multiple GSC isolates and found one additional GSC isolate that showed requirement for *FBXO42* (Figure 1B)(Table S1). Given this seemingly low percentage, we next explored requirement among a broad sampling of cancer cell lines using data now available through the Broad and Sanger Institutes [23, 24]. This included the use of cell line gene effect scores or CERES scores from CRISPR-Cas9 where *FBXO42* was included as a target [23] (Figure 1C). Using a cutoff of -0.5, we observed that cancer cell lines from most tissue lineages contain a subset of *FBXO42* loss-sensitive (F42L-*S*) cell lines, with a range of 0-30%, a mean of 13.2%, and a median of 11.1% (Figure 1C). Across all tissues, approximately 15% of cell lines were predicted to be potentially F42L-*S*, in-line with the frequency we observed for GSC isolates.

To substantiate these finding further, we performed individual retests using an optimized method for nucleofection of sgRNA:Cas9 ribonucleoprotein complexes [25]. We nucleofected GSC-0827 and NSC-U5 with CRISPR RNPs targeting *FBXO42, HEATR1* (an essential gene control) and *CD8A* (a non-essential, gene targeting control) and measured relative cell viability. We found that loss of *FBXO42* did indeed have a profound effect on F42L-*S* GSC-0827 cells, with almost no cells surviving, without affecting predicted F42L-*R* GSC-0131 and NSC-U5 cells (Figure 1D). Importantly, we observed a strong reduction in viability in NSC-U5 when targeting *HEATR1*, indicating that our differential results were not due to reduced nucleofection efficiency in GSC-0131 and NSC-U5 cells.

We also examined the predicted F42L-*S* cell lines from Figure 1C. We picked four F42L-*S* cell lines – the breast cancer ductal carcinoma line BT549, the osteosarcoma line 143B, the glioblastoma line T98G, and the gastric adenosquamous carcinoma line MKN1 – and nucleofected RNPs as above. These four lines showed exquisite (∼ equal to *HEATR1* loss) or moderate (significant but less than *HEATR1* loss) F42L sensitivity. Taken together, the results suggest the F42L-*S* phenotype is applicable to significant subset of cancers arising from a broad range of tissues.

### *FBXO42* inactivation compromises F42L-*S* GSC tumor growth

Next, we wished to determine if the F42L-*S* phenotype holds true *in vivo* in GSC-derived tumors. We first engineered GSC-0827 cells such that using Doxycycline (Dox) controllable ectopic expression of *FBXO42* to complement disruption endogenous *FBXO42* locus (Methods). We identified outgrown clones that recapitulated loss of *FBXO42* in the absence of Dox. Clone 15 contained a net frameshift mutation in each allele (Figure S1) and loss of FBXO42 protein expression after 4 days of Dox removal (Figures 1F). To ensure that Dox would be bioavailable to tumor cells, we used a flank tumor model to evaluate effect of *FBXO42* loss after tumors had already formed. We observe that after attenuating *FBXO42* expression via Dox removal, GSC-0827 tumors show a substantial loss in tumor volume, almost becoming undetectable, before gradually regrowing. The results suggest that *FBXO42* is requirement for maintenance of tumor growth in F42L-*S* cells.

### *FBXO42*’s substrate recognition and SCF-interaction domains are required for viability in F42L-*S* cells but not its known substrates

Since FBXO42 is an F-box containing protein that serves as the substrate-recognition component of an SCF (SKP1-CUL1-F-box protein)-type E3 ubiquitin ligase complex [26], we hypothesized that failure to degrade one or more of its substrates a key to F42L-*S* phenotype. Knowledge of the functional domains of FBXO42 allowed us to test the general hypothesis that its participation in an E3 ligase complex that causes F42L-*S*.

F-box proteins interact with SKP1 through their F-box domain, while they interact with ubiquitination targets through other protein interaction domains. Besides an F-box domain, FBXO42 additionally contains solely a Kelch repeat domain [27], which serves as its substrate-binding domain [19, 20, 22, 27]. Therefore, we created lentiviral constructs containing 3XFLAG-tagged ΔF-box domain (deletion of aa 39-93) and ΔKelch domain (deletion of aa 107-354) mutant versions of *FBXO42*, along with a 3XFLAG-tagged full-length *FBXO42* control construct. We then transduced GSC-0827 cells with these constructs, which also conferred puromycin resistance, and selected them with puromycin. In order to test whether exogenous expression of the full-length, ΔF-box, and ΔKelch versions was able to rescue loss of endogenous *FBXO42*, we nucleofected the cells with CRISPR RNPs for two *FBXO42* sgRNAs to which the lentiviral vectors had been designed to be resistant (via synonymous mutation of multiple bases and use of an sgRNA spanning an intron-exon boundary), along with non-targeting and targeting control sgRNAs, and subsequently measured relative cell viability (Figure 2A). We observed that relative to the sgCD8A targeting controls, expression of full-length *FBXO42* was able to almost fully rescue the viability loss observed upon endogenous *FBXO42* knockout. However, with expression of the ΔF-box and ΔKelch versions, we observed a similar viability loss upon *FBXO42* knockout as we observed in untransduced control GSC-0827 cells (Figure 2A). Importantly, the protein expression levels of the ΔF-box and ΔKelch versions were similar to expression of the full-length version (Figure 2B), indicating that the difference was not simply due to insufficient expression of the ΔF-box and ΔKelch proteins. These data suggest that both the F-box domain and the Kelch domain are required for survival of F42L-*S* cells, which supports the idea that FBXO42’s role in an E3 ubiquitin ligase complex is responsible for the observed phenotype.

**Figure 2:**
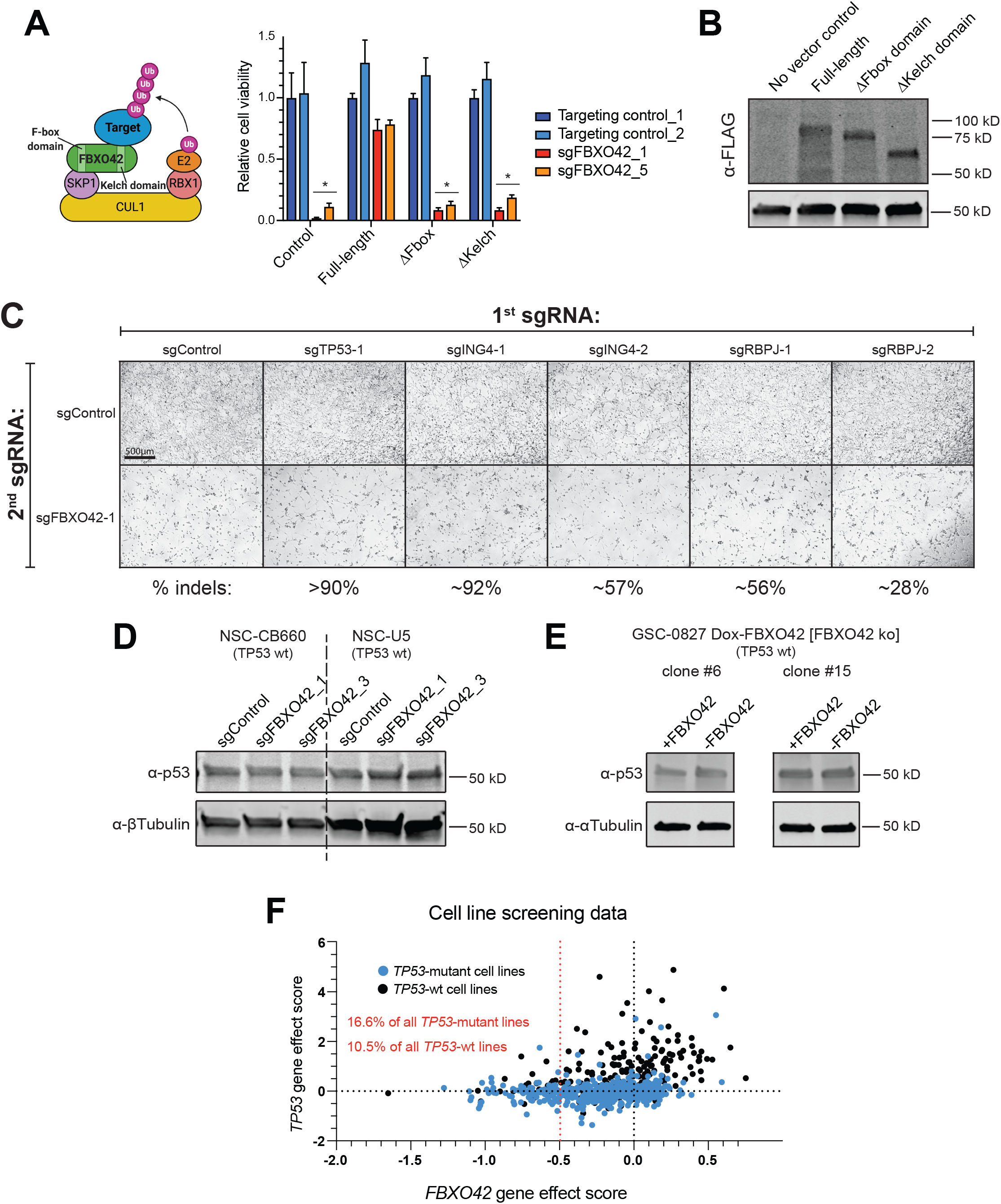
*FBXO42*’s ubiquitin ligase activity-associated domains are required in FBXO42 sensitivity cells, but not its known targets. **A**, Relative cell viability (normalized to targeting control *sgCD8A*) for GSC-0827 transduced with full-length, F-box domain deletion mutant, or Kelch domain deletion mutant versions of 3XFLAG-*FBXO42* and then nucleofected with CRISPR RNPs targeting *FBXO42*, measured at 8 days post nucleofection. The lentiviral expression constructs are resistant to the *FBXO42* sgRNAs used here. Untransduced GSC-0827 are shown for comparison. sgNTC = non-targeting control sgRNA. (n=3; *pval.<.01 Student’s t-test) **B**, Western blot for FLAG tag in transduced GSC-0827 used for viability assay in **A**. **C**, Representative images of GSC-0827 nucleofected with CRISPR RNPs targeting the published FBXO42 targets/interactors TP53, ING4, or RBPJ in combination with *sgNTC* or *sgFBXO42*, taken at 5 days post nucleofection. **D, E**, Western blots for p53 levels in **D**, NSCs nucleofected with CRISPR RNPs targeting *FBXO42* and **E**, GSC-0827 clones 6 and 15 4 days after Dox removal. **F**, Cell line functional genomic screening data from DepMap showing *FBXO42* dependency score vs. *TP53* dependency score (CERES scores from CRISPR Avana 19Q4 dataset). *TP53*-mutant cell lines are marked in blue.

Building upon this idea, we next wanted to know if a known target of FBXO42 is responsible for the viability phenotype. In this scenario, knocking out the target should rescue the viability loss seen with *FBXO42* knockout in F42L-*S* cells. There are three published targets of FBXO42, including TP53 [19, 20], ING4 [22] (a chromatin reader), and RBPJ [28] (a transcriptional regulator important in Notch signaling). We performed knockout rescue experiments, whereby RNPs containing *FBXO42* or control sgRNAs were co-nucleofected with sgRNAs targeting *TP53, ING4*, and *RBPJ*. Co-nucleofections do not diminish the effect of each targeting in our conditions [25]. However, as shown from representative micrographs shown in Figure 1C, knockout of these targets where not essential in GSC-0827 cells and they also failed to rescue the F42L-*S* phenotype.

Of these targets, one group reported that FBXO42 directly interacts with p53 to cause its ubiquitination and degradation. They argued that loss of FBXO42 causes p53 stabilization, leading to G1 cell cycle arrest and apoptosis [19, 20]. Because of the importance of this possibility, we wanted to further test whether FBXO42 could be regulating p53 in p53 wt F42L-*R* and *-S* cells. We assessed steady-state p53 protein levels in NSCs (p53 wt), which we had nucleofected with CRISPR RNPs targeting *FBXO42*, as well as in our dox-*FBXO42* inducible clones 6 and 15 of F42L-*S* GSC-0827 cells (Figure 2D & E). However, we did not observe any increase in steady-state levels of p53.

Furthermore, we reasoned that if the negative effect on viability observed with *FBXO42* loss were indeed due to stabilization of p53, then *TP53*-mutant cancer cell lines should be less likely to be sensitive to *FBXO42* loss than *TP53*-wt cell lines. To address this, we examined the relationship between *FBXO42* dependency, *TP53* dependency, and *TP53* mutation status among cancer cell lines from Figure 1C (Figure 2F). With regard to *TP53* status versus *TP53* gene effect score, the non-mutant cell lines tend to show positive effect scores due to enhancement of proliferation. However, the results clearly show that *FBOX42* requirement does not depend on *TP53* status, where a similar percentage of F42L-*S* cells are found in both categories. If *FBXO42* requirement were dependent on p53 status (e.g., similar to MDM2), there should be a strong negative trend among TP53 wt cell lines, which is not at all apparent (Figure 2F).

Taken together, our data suggest that FBXO42’s role in SCF is responsible for its viability requirement in F42L-*S* cells, but that the effect is likely mediated through one or more, as yet unidentified substrates. Moreover, the results demonstrate that p53 is not a significant target of FBXO42.

### CCDC6 requirement is associated with FBXO42 requirement in F42L-*S* cells

To identify candidate protein interactors of FBXO42, we examined protein-protein interactions found in large-scale mass-spec based databases, which incorporated FBXO42 [29, 30]. We found several known interactors, including CUL1, SKP1, and RBPJ, and a few novel candidates, including CCDC6, PPP4C and PPP4R1 (Figure 3A). To determine relevance, we examined requirement for these among cancer cell lines showing requirement for *FBXO42*. Only one showed a significant correlation with *FBXO42* requirement: *CCDC6* (Figure 2B).

**Figure 3:**
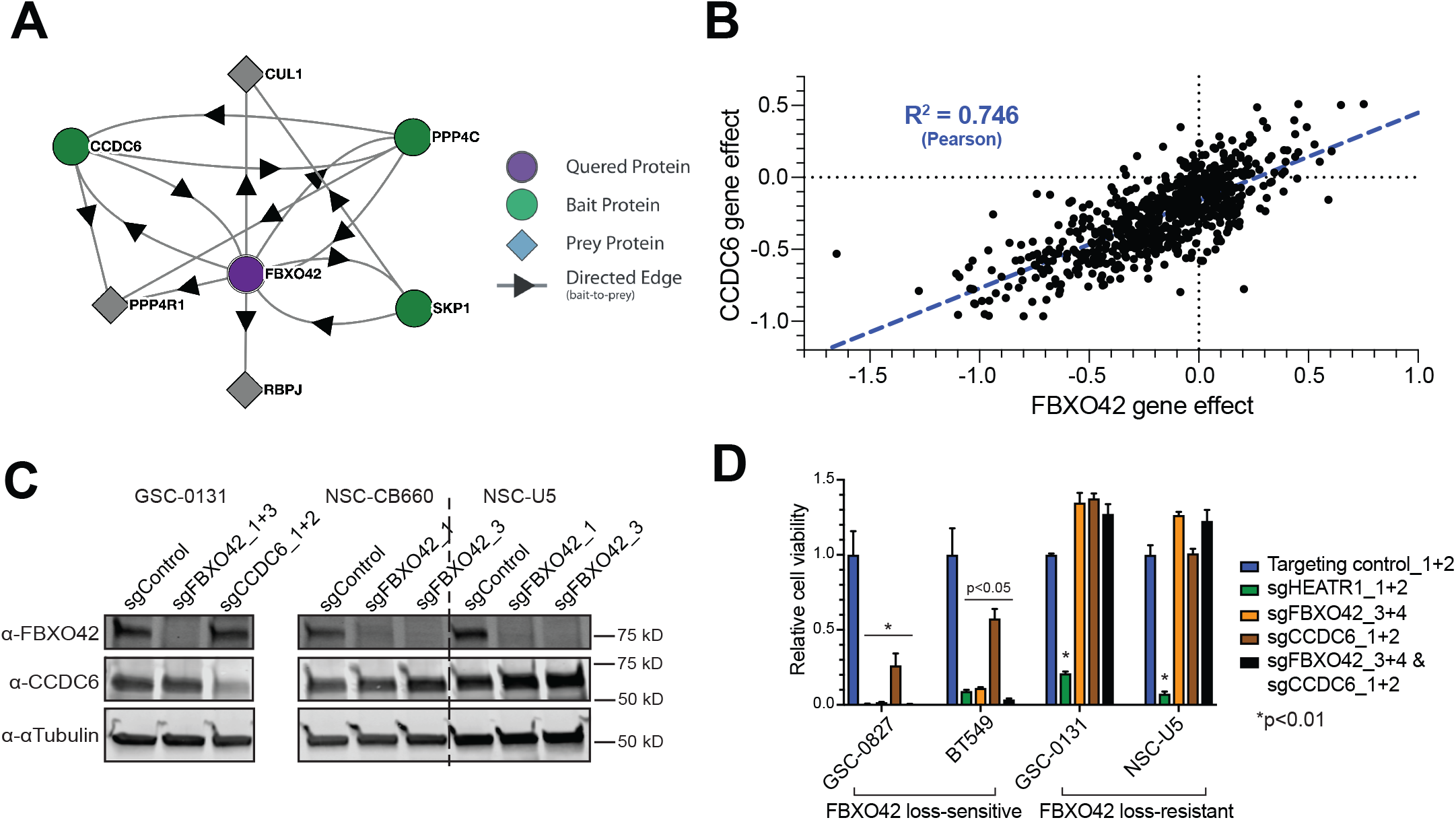
*CCDC6* requirement is associated with *FBXO42* requirement in F42L-*S* cells but is not a degradation target of FBXO42. **A**, Protein-protein interactions for FBXO42 predicted by BioPlex (PMID: 33961781). **B**, Cell line functional genomic screening data from DepMap showing *FBXO42* dependency score vs. *CCDC6* dependency score (CERES scores from CRISPR Avana 19Q4 dataset). Each dot represents a cell line. Dotted blue line shows linear regression fit. **C**, Western blot for FBXO42 and CCDC6 levels in GSCs and NSCs nucleofected with CRISPR RNPs targeting *FBXO42* or *CCDC6*, 4 days post-nucleofection. **D, Relative** cell viability (normalized to targeting control *sgCD8A*) for cells nucleofected with CRISPR RNPs targeting *FBXO42, CCDC6*, or both. *HEATR1* is an essential control gene. *sgNTC* = non-targeting control sgRNA. Measured at 8-11 days post nucleofection (depending on doubling time).

CCDC6 is a coiled-coil domain-containing protein that is considered a tumor suppressor and was first identified due to its involvement in chromosomal rearrangements with the RET proto-oncogene in thyroid papillary carcinomas [31]. It is a pro-apoptotic protein substrate of ATM that has been shown to be involved in the DNA damage response [32-34]. However, *FBXO42* KO did not affect steady-state levels of CCDC6 (Figure 3C), suggesting it is not a target of FBXO42.

We next assessed relative cell viability after nucleofecting two F42L-*S* lines GSC-0827 and BT549 and the two F42L-*R* lines with CRISPR RNPs targeting *FBXO42* or *CCDC6* individually or both in combination (Figure 3D). The results confirmed that *CCDC6* dependency is associated with *FBXO42* dependency in our system as well, since *CCDC6* knockout reduced viability in GSC-0827 and BT549 but had no effect in GSC-0131 or NSC-U5. Combined loss of *FBXO42* and *CCDC6* may have a greater impact on viability in F42L-*S* cells than loss of either gene alone; however, due to the limitations of the assay and viability effects, epistasis seems just as likely (Figure 3D). For F42L-*R* cells, single or combined loss of either gene did not affect their outgrowth (Figure 3D). These results indicate that F42L-*S* cells require both genes for survival, that *FBXO42* and *CCDC6* are not synthetic lethal, and that CCDC6 likely participates in FBXO42 function in F42L-*S* cells.

### *FBXO42* loss triggers an extended mitotic arrest with misaligned chromosomes in F42L-*S* cells

We next wanted to gain insight into the cause of *FBXO42* dependency in F42L-*S* cells. Since many E3 ubiquitin ligases are involved in cell cycle control, we first determined changes in gene expression. If cells were arrested in a specific part of the cell cycle, we would expect to see a dramatic increase in cell cycle genes peaking in that phase (e.g., [35]). Therefore, we examined gene expression changes induced by *FBXO42* loss in F42L-*S* and F42L-*R* cells. Figure 4A shows that in F42L-*R* NSCs only 5 genes showed significant differences, while in F42L-*S* GSC-0827 cells display significant changes in ∼523 genes. For the 399 gene upregulated, there was significant enrichment for genes whose expression peaks in G2/M, including those involved in chromosome segregation and mitosis, such as *BUB1B, CCNB1, CDC20, CENPE, GTSE1*, and *PLK1* [36](Figure 4B)(Table S3).

**Figure 4:**
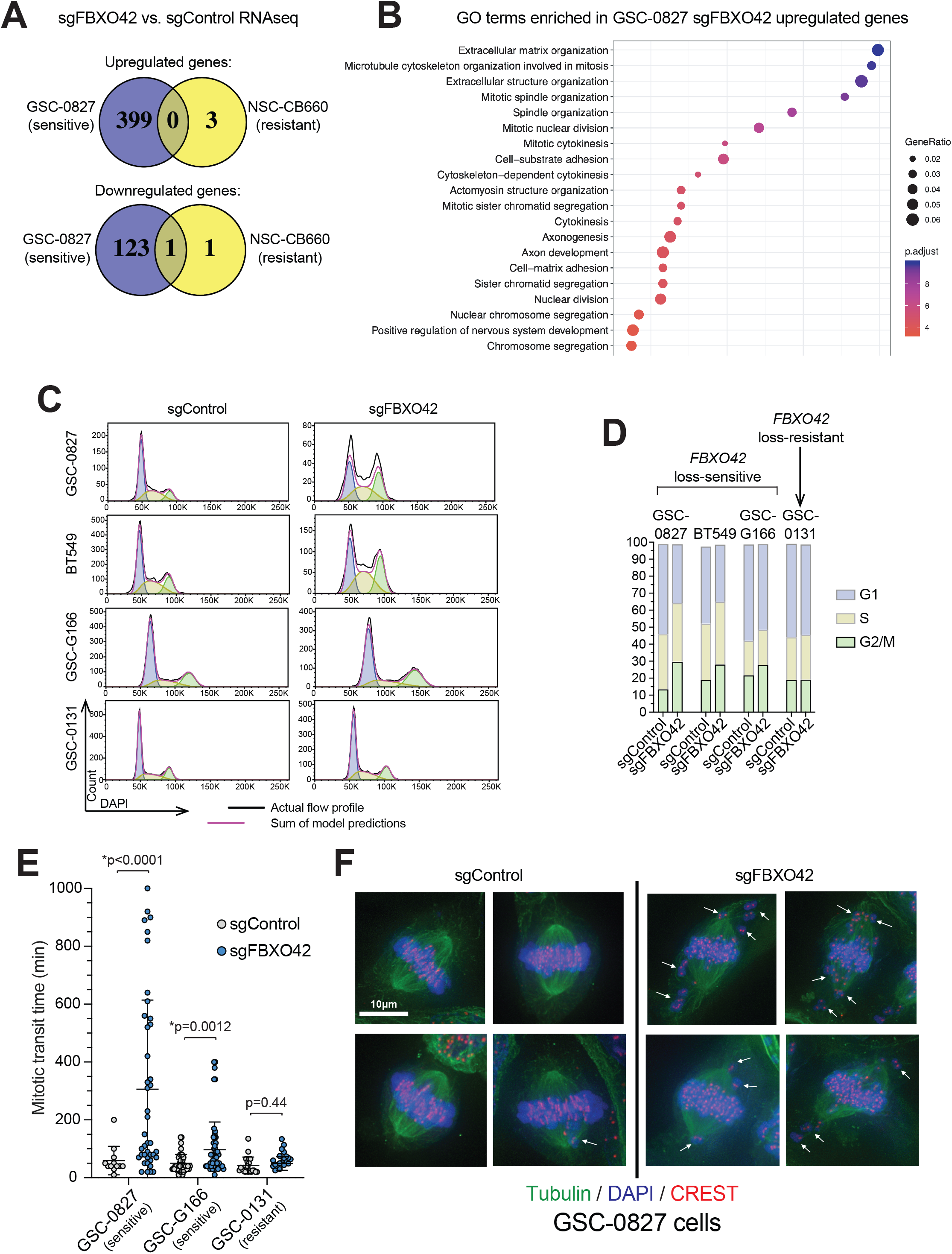
*FBXO42* loss triggers an extended mitotic arrest associated with misalignment of chromosomes in metaphase in F42L-*S* cells. **A**, Comparison of gene expression changes after knockout of *FBXO42* in GSC-0827 and NSC-CB660 cells (n=3; 4 days post-nucleofection). **B**, Gene set enrichment analysis of genes up regulated after *FBXO42* knockout in GSC-0827 cells. **C**, DNA content (DAPI) flow cytometry profiles for cells nucleofected with CRISPR RNPs targeting *FBXO42* compared to a non-targeting sgRNA (*sgNTC*) (4 days post nucleofection). Dean Jett Fox model for cell cycle distribution (FlowJo software) is shown under the histogram for each sample, with predictions for G1, S, and G2/M in violet, yellow, and green, respectively. **D**, Cell cycle proportion of DNA content (DAPI) flow cytometry profiles for cells nucleofected with CRISPR RNPs targeting *FBXO42* compared to a non-targeting sgRNA (*sgNTC*) from **C**. **E**, Mitotic transit time, measured using analysis of time-lapse microscopy, for individual H2B-EGFP-expressing cells with or without knockout of *FBXO42*. Bars show mean and standard deviation. *Indicates significance as shown, student’s t-test. **F**, Chromosome alignment assays in *FBXO42* knockout and control GSC-0827 cells 4 days post-nucleofection were treated by 10 μM MG-132 for 2 hours to arrest them at metaphase and then fixed, stained as indicated (CREST anti-serum stains human kinetochores) and visualized using deconvolution microscopy. Scale bar indicates 10 microns.

These results suggested that *FBXO42* loss in F42L-*S* cells triggers a G2/M or M-phase cell cycle arrest. We, thereby, assessed the cell cycle profiles of F42L-*S* and F42L-*R* cells with and without *FBXO42* knockout using flow cytometry DNA content analysis via DAPI staining (Figure 4C). There was a marked increase in the percent of cells in the G2/M stage in F42L-*S* cells treated with *sgFBXO42*, with a concomitant decrease in the percent in G1 (Figure 4C & D). The degree of F42L-*sensitivity* was also associated with the increase with the present of cells observed in G2/M, as the exquisitely sensitive lines GSC-0827 and BT549 displayed the largest increase while the moderately sensitive line GSC-G166 displayed a lesser increase. Importantly, there was no difference in cell cycle profile upon *FBXO42* loss in the F42L-resistant line GSC-0131 (Figure 4D).

Since it is not possible to distinguish between cells in G2 and M phase when using a DAPI DNA content profile to assess cell cycle, we used time-lapse microscopy in GSCs that had been transduced with a PGK-H2B-EGFP construct to determine if we could see an effect on mitosis in *FBXO42* knockout cells (Figure 4E). We imaged cells (phase and GFP) every 5 minutes to compile time-lapse videos and then assessed the mitotic transit time for individual cells. We found that F42L-*S* cells with *FBXO42* knockout spent a significantly longer time in mitosis. Once again, the difference was associated with the degree of F42L-sensitivity, with GSC-0827 sgFBXO42 cells spending an average of ∼3.1 times as long in mitosis as control cells (up to a maximum of ∼8.9 times) and GSC-G166 *sgFBXO42* cells spending an average of ∼1.50 times as long in mitosis as control cells (up to a maximum of ∼4.9 times) (Figure 4E). We also observed that GSC-0827 cells failed to adapt to their mitotic arrest and eventually suffered cell death during this arrest, while GSC-G166 cells were eventually able to overcome the arrest and exit mitosis. As expected, based on the flow cytometry results, there was no significant difference in mitotic transit time between *sgFBXO42* and control cells for the F42L-*R* isolate GSC-0131 (Figure 4E).

We observed from our time-lapse microscopy that the mitotic arrest seemed to be specifically a metaphase arrest, since we could see a formed metaphase plate in the arrested cells (visualized by H2B-EGFP). We wanted to explore this further and thus fixed *sgFBXO42* and control GSC-0827 cells, stained with DAPI and tubulin and centromere specific antibodies, and performed Z-stack ultra-high-resolution microscopy to create 3D projections of mitotic cells (Figure 4F). This confirmed that arrested cells were indeed at metaphase, and it showed that *sgFBXO42* cells suffer from both a distorted spindle and a dramatic increase in chromosome alignment defects, with many misaligned chromosomes present at the spindle poles (Figure 4F).

Altogether, these results indicate that loss of *FBXO42* in F42L-*S* cells leads to a prolonged mitosis, likely due to defects in chromosome congression and alignment.

### Inhibition of spindle assembly checkpoint kinase Mps1 suppresses mitotic arrest, but not loss viability loss, triggered by *FBXO42* inactivation in F42L-*S* cells

A metaphase arrest triggered by misaligned chromosomes would be predicted to activate of the spindle assembly checkpoint (SAC), which is a feedback-control system in eukaryotic cells that monitors the attachment of kinetochores to the microtubule fibers of the mitotic spindle [37]. Sister chromatids that do not have proper bi-oriented attachments at kinetochores cause SAC signaling, activating the SAC effector, the mitotic checkpoint complex (MCC) [37]. The MCC binds and inhibits APC/C-Cdc20, which is required for the metaphase–anaphase transition, thus, preventing entry into anaphase. In this manner, the SAC serves to prevent premature chromosome segregation in the presence of chromosomes that are not properly attached to the spindle, thereby preserving the genome from the disastrous consequences that aneuploidy can bring [38]. The activity of the Mps1 kinase is required to activate the SAC by phosphorylating the kinetochore protein Knl1 [39-41], creating docking sites for the recruitment of additional SAC proteins [42-48]. Its chemical inhibition prevents activation of the SAC.

To determine if SAC activation was the cause of the prolonged metaphase arrest in F42L-*S* cells, we asked whether a brief treatment with an Mps1 inhibitors alleviates the mitotic arrest triggered by loss of *FBXO42*. To this end, we used the Dox-inducible *FBXO42* GSC-0827 cells (clone 15) described above. We cultured cells with or without doxycycline for 4 days, a timepoint which corresponded to complete loss of FBXO42 protein and the beginning of the mitotic arrest phenotype. We then treated these cells with either Mps1 inhibitor or vehicle control for 2 hours and performed flow cytometry for DAPI and p-H3 (Ser10) to mark mitotic cells. From the vehicle controls, we could see that at this timepoint >30% of cells in the Dox-condition (*FBXO42* loss) were arrested mitosis, compared to steady-state level of 2.25% in the Dox+ control. After treatment with Mps1 inhibition only 3.8% of cells were in mitosis in the Dox-condition, indicating that Mps1 activity is required for the mitotic arrest phenotype.

We also assessed the relative cell viability in cells cultured with or without Dox and with or without Mps1 inhibition (Figure 5C). Although there was a partial rescue of viability in the presence of the inhibitor, overall, the cells still lost viability after treatment (Figure 5C). This likely indicates that chromosome segregation errors caused by loss of *FBXO42* are of sufficient magnitude to cause significant loss of viability, regardless of SAC activation. We conclude that the mechanism of cancer-specific requirement of *FBXO42* is to promote congression and kinetochore-microtubule attachment and prevent triggering of the SAC.

**Figure 5:**
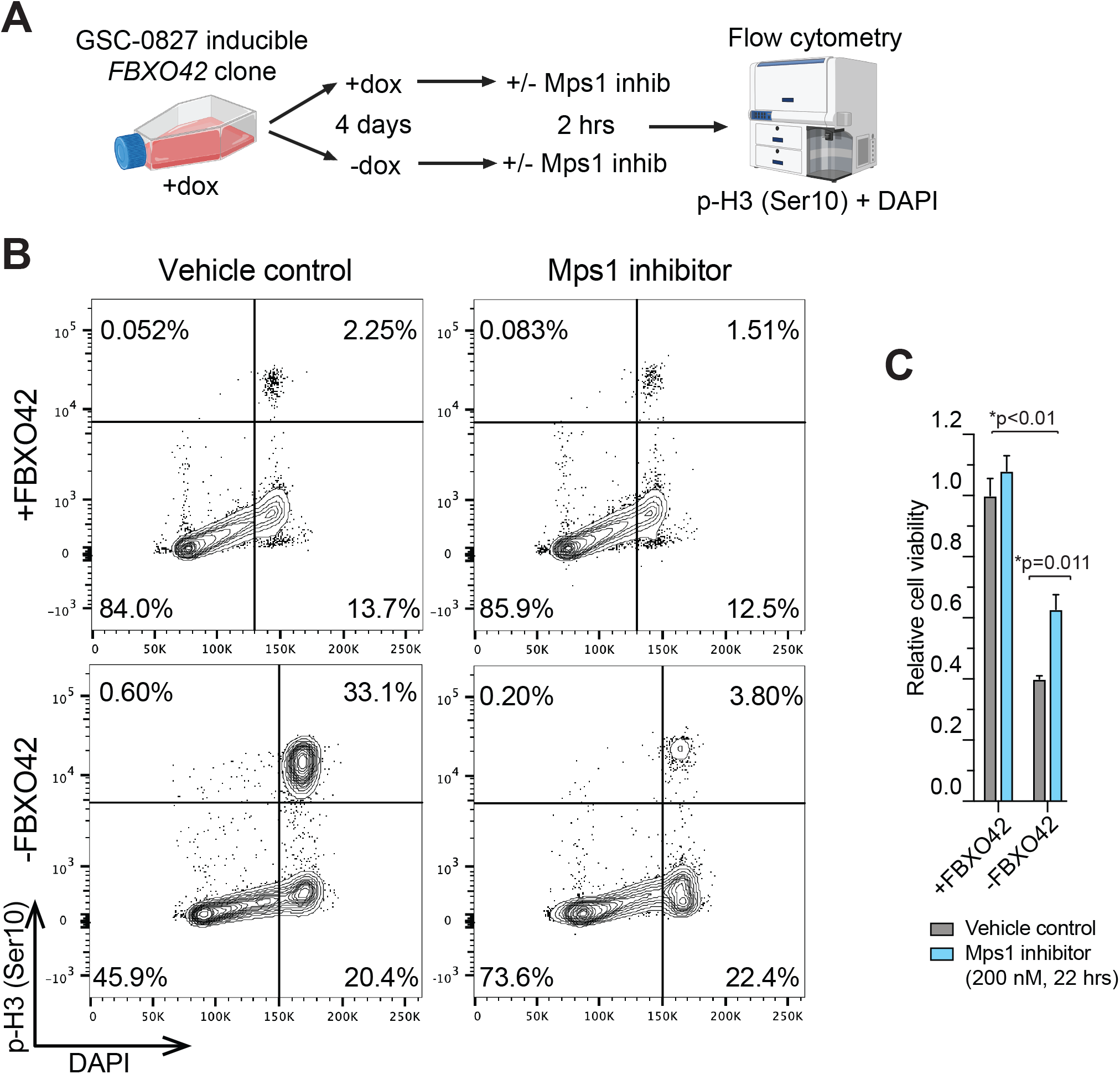
Inhibition of spindle assembly checkpoint kinase Mps1 by-passes G2/M arrest, but not loss of viability, triggered by *FBXO42* inactivation in F42L-*S* cells. **A**, Overview of experiment shown in B. **B**, DAPI vs. p-H3 (Ser10) flow cytometry profiles for GSC-0827 doxycycline-inducible *FBXO42* clone 15 kept in +/- doxycycline for 4 days and then treated with vehicle or an Mps1 inhibitor (NMS-P715) for 2 hours. **C**, Relative viability for cells that were kept in +/- doxycycline for 4 days and then treated with vehicle or 200 nM Mps1 inhibitor for 22 hours.

## Discussion

Here, we characterized cancer-specific requirement for the F-box protein encoding gene *FBXO42*. Starting with GBM isolates we found a novel viability requirement in ∼15% of cancer cell lines that seems independent of tissue of origin (Figure 1), which was also observed in patient-derived xenograft tumors. In F42L-*S* cells *FBXO42* was required to prevent mitotic delay or extended arrest driven by SAC activation. We observed that FBXO42’s F-box and Kelch domains were critical to maintaining viability of F42L-*S* cells, implicating its SCF-associated ubiquitin ligase activity. However, none of FBXO42’s known substrates were required, as deleting them had no effect on *FBXO42* loss phenotype. In contrast to two previous reports, we also show that FBXO42 does not affect steady state p53 levels or modulate cell growth of p53 wt in a manner consistent with p53 being a degradation target. Instead, we propose a model of *FBXO42* requirement in which FBXO42 degrades a novel cancer-specific target in F42L-*S* cells (Figure 6). In the absence of FBXO42 activity, the presumed degradation target accumulates and perturbs chromosome-spindle dynamics, triggering spindle assembly checking arrest and ultimately cell death.

**Figure 6:**
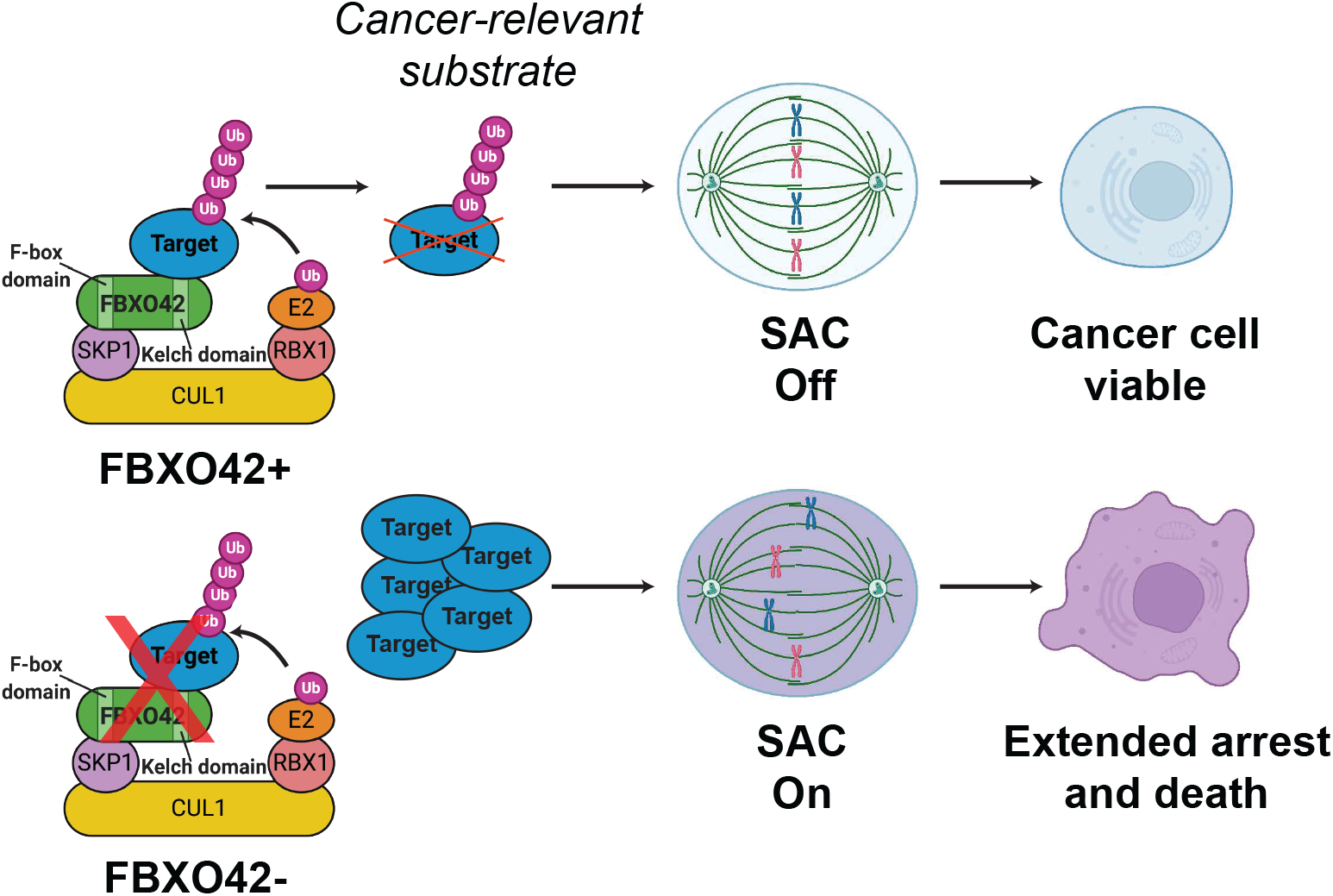
A model for cancer-specific requirement of *FBXO42*.

A recent study examining roles for E3 ubiquitin ligase complex genes in broad range of biological processes, including cell cycle modulatory drugs, identified *FBXO42* mutant cells as being sensitive to mitotic inhibitors, including BI-2536 (Plk1) and colchicine (microtubules)[49]. They were further able to show that in *FBXO42* mutant HAP1 cells, there was an increased frequency of monopolar spindles upon BI-2536. While they were not able to establish a mechanism, the results suggest that FBXO42 activity helps buffer the effects of perturbations in chromosome-spindle dynamics. Given these results along with our own, it is likely that *FBXO42* requirement in F42L-*S* cells is caused by an underlying perturbation in chromosome-spindle dynamics.

Consistent with this notion, we observed chromosome alignment defects in sensitive GSC-0827 cells, which is the classic cause of SAC activation (reviewed in[50]). In addition, we also observed that the mitotic spindle was noticeably twisted and distorted (Figure 4F). Therefore, it is also possible that FBXO42 substrate(s) act on the mitotic spindle and, thereby, causes prolonged activation of the SAC [51].

For GBM isolates, we have previously established that there are at least two cancer-specific defects in kinetochore-microtubule dynamics that can cause profound loss of chromosome-spindle attachments [37]. The most dramatic of these is caused by inappropriate Ras/MAP kinase activity in mitosis [12, 13, 18]. About half of GBM isolates tested suffer from this defect, which causes a non-essential domain of BUB1B/BubR1 (its GLEBs domain) to become essential [12, 18]. However, F42L-*S* GSC-0827 cells do not have this kinetochore defect [18]. Thus, *FBXO42* requirement represents a separate and novel form of mitotic perturbation. Identifying its key mitotic-relevant substrate(s) will likely be key to understanding the mechanism of its cancer-specific requirement.

However, we were unable to identify FBXO42’s relevant substrate(s) for the phenotypes reported above, either from those published or ones we attempted to identify by IP-Mass spec (data not shown). We were able to confirm requirement for CCDC6, which appears in protein-protein interaction databases as interacting with FBXO42. However, it is unlikely to be a substrate of FBXO42. Other candidate interacting proteins found in these databases include members of the PPP4C-PPP4R1 PP4 phosphatase (Figure 2A). This complex can affect microtubule organization via negative regulation of CDK1 interphase, which intern causes abnormal phosphorylation of NDEL1 [52], which localizes to kinetochores, affects chromosome alignment, and triggers SAC when inhibited [53]. Future experiments will be required to examine these and other possibilities regarding *FBXO42* function.

## Acknowledgements

We thank members of the Paddison and Olson Lab for helpful discussions, and Pam Lindberg and An Tyrrell for administrative support. This work was supported by the following grants: Interdisciplinary Training in Cancer Fellowship NCI T32CA080416 (P.H.); NCI/NIH (R01CA190957) (P.P.) (R01CA114567) (J.O.); and NINDS/NIH (R01NS119650) (P.P.). This research was funded in part through the NIH/NCI Cancer Center Support Grant P30 CA015704.

## Author Contributions

Project conception and design was carried out by P.H. and P.P.; experiments and data analysis were performed by P.H., E.G., and J.H.; critical reagents were generated by M.K.; bioinformatical data analysis and statistics were performed by P.H., S.A., and P.P.; P.P. and P.H. wrote the manuscript with input from other authors. Funding acquisition: J.O. and P.P.

## Methods

*Key Reagents and Resources are available in Table S3*.

### Cell Culture

Isolates were cultured in NeuroCult NS-A basal medium (StemCell Technologies) supplemented with B27 (Thermo Fisher Scientific), N2 (homemade 2x stock in Advanced DMEM/F-12 (Thermo Fisher Scientific)), EGF and FGF-2 (20 ng/ml) (PeproTech), glutamax (Thermo Fisher Scientific), and antibiotic-antimycotic (Thermo Fisher Scientific). Cells were cultured on laminin (Trevigen or in-house-purified)-coated polystyrene plates and passaged as previously described [54], using Accutase (EMD Millipore) to detach cells.

### Lentiviral Production

For virus production, lentiCRISPR v2 plasmids [55] were transfected using polyethylenimine (Polysciences) into 293T cells along with psPAX and pMD2.G packaging plasmids (Addgene) to produce lentivirus. For the whole-genome CRISPR-Cas9 libraries, 25×150mm plates of 293T cells were seeded at ∼15 million cells per plate. Fresh media was added 24 hours later and viral supernatant harvested 24 and 48 hours after that. For screening, virus was concentrated 1000x following ultracentrifugation at 6800*xg* for 20 hours. For validation, lentivirus was used unconcentrated at an MOI<1.

### CRISPR-Cas9 Screening

For screening, cells were transduced to achieve ∼750X representation of the library (at ∼30% infection efficiency to ensure a high proportion of single integrants). 2 days after transduction, media was replaced with media containing 2 μg/mL puromycin. After 3 days of selection, portions of cells representing 500-750X coverage of the library were collected as the “Day_0” samples. The remaining cells were cultured and consistently maintained at 500-750X representation for 21-23 days, after which time the “Day final” samples were collected. Screening was carried out in triplicate. To read out screen results, genomic DNA was extracted using the QIAamp DNA Blood Mini Kit (QIAGEN), and a two-step PCR procedure was used to first amplify the genomically integrated sgRNA sequences and then to incorporate Illumina deep sequencing adapters and barcodes onto the sgRNA amplicons. For the first round of PCR, a sufficient number of PCR reactions were carried out to use all gDNA from the 500-750X coverage sample of cells at 2μg genomic DNA per PCR reaction, using MagniTaq Multiplex PCR Master Mix (Affymetrix) and 12 cycles. For the second round of PCR, 5μL of the first-round product was used as a template in combination with primers that would add the deep sequencing adapters and barcodes, using Herculase II Fusion DNA Polymerase (Agilent) and 16 cycles. Amplicons from the second round PCR were then column purified using the PureLink Quick PCR Purification Kit (Invitrogen). Purified PCR products were sequenced using HiSeq 2500 (Illumina). Bowtie [56] was used to align the sequenced reads to the sgRNA library, allowing for 1 mismatch. The R/Bioconductor package edgeR [57] was used to assess changes across groups.

### RNA-seq analysis

Cells were lysed with Trizol (Thermo Fisher). Total RNA was isolated (Direct-zol RNA kit, Zymo Research) and quality validated on the Agilent 2200 TapeStation. Illumina sequencing libraries were generated with the KAPA Biosystems Stranded RNA-Seq Kit[58] and sequenced using HiSeq 2000 (Illumina) with 100bp paired-end reads. RNA-seq reads were aligned to the UCSC hg19 assembly using STAR2 (v 2.6.1)[59] and counted for gene associations against the UCSC genes database with HTSeq [60]. Normalized gene count data was used for subsequent hierarchical clustering (R package ggplot2 [61]) and differential gene expression analysis (R/Bioconductor package edgeR [57]). Heatmaps were made using R package pheatmap [62].

### Western Blotting

Cells were harvested, washed with PBS, and lysed with modified RIPA buffer (150mM NaCl, 25mM Tris-HCl (pH 8.0), 1mM EDTA, 1.0% Igepal CA-630 (NP-40), 0.5% sodium deoxycholate, 0.1% SDS, 1X protease inhibitor cocktail (complete Mini EDTA-free, Roche)). Lysates were sonicated (Bioruptor, Diagenode) and then quantified using Pierce BCA assay (Thermo Fisher). Identical amounts of proteins (20-40μg) were electrophoresed on 4–15% Mini-PROTEAN TGX precast protein gels (Bio-Rad). For transfer, the Trans-Blot Turbo transfer system (Bio-Rad) with nitrocellulose membranes was used according to the manufacturer’s instructions. TBS (137mM NaCl, 20mM Tris, pH 7.6) +5% nonfat milk was used for blocking, and TBS+0.1%Tween-20+5% milk was used for antibody incubations. The following commercial primary antibodies were used: MX1 (Cell Signaling #37849S, 1:500), Tp53 (Cell Signaling #48818, 1:500), αTubulin (Sigma #T9026, 1:1,000), ΔActin (Cell Signaling #3700S, 1:2,000). The following secondary antibodies were used (LI-COR, all 1:10,000): #926-68073, #926-32212, #926-32214, #926-68074, #925-32212, #925-68071. An Odyssey infrared imaging system (LI-COR) was used to visualize blots.

### Flow Cytometry

Processed cells were flow cytometry analyzed immediately using either a BD FACSymphony A5 or BD LSRFortessa X-50 machine. Results were analyzed using FlowJo software.

### Viability Assays

Viable cell numbers were measured using CellTiter-Glo Luminescent Cell Viability Assay (Promega) according to manufacturer’s instructions.

### Time-lapse Microscopy

Cells were first infected with lentivirus for PGK-H2B-EGFP (Addgene) and then nucleofected with CRISPR RNPs as needed and plated into 24-well or 12-well plates. 1-3 days post nucleofection (depending on cell line and onset of gene knockout phenotype), cells were placed into an IncuCyte S3 (Sartorius) instrument. For the mitotic transit time analysis, phase and fluorescence (GFP) images were taken every 5 minutes for 48-72 hours. Videos were compiled using the IncuCyte S3 software, and mitotic transit time was then analyzed for individual cells. A cell was considered to enter mitosis when nuclear envelope breakdown was evident based on EGFP visualization and when a visible morphology change from flat to round was observed. Following successful cytokinesis (proper cytoplasmic division resulting in two daughter cells), a cell was categorized as having successfully completed mitosis. A cell was classified as cytokinesis failure if the cell failed to divide following mitotic entry due to an abrupt mitotic exit while in metaphase or anaphase, or failure to complete cytokinesis. If a cell seemed to experience cytokinesis failure, it was followed for additional time to ensure that this was indeed the case. A cell was categorized as cell death in mitosis if a cell erupted and died during mitosis (between nuclear envelope breakdown and cytokinesis).

### Generation of Dox-inducible FBXO42 cells

We cloned an *sgFBXO42_1*-resistant version of the *FBXO42* ORF into the retroviral Tet-On vector pTURN-tight and transduced GSC-0827 cells with virus for this construct. Keeping the cells on doxycycline to maintain exogenous *FBXO42* expression, we then nucleofected them with CRISPR RNPs for sgFBXO42_1 in order to knock out the endogenous *FBXO42* alleles. To obtain cells with a uniform and well-controllable induction of exogenous expression, we then derived clones from this cell pool. We screened the clones for proper ability to turn off the construct by taking a subset of cells of each clone and testing for viability loss upon doxycycline removal. Based on these results, we took our top two clones (clones 6 and 15), PCR amplified the endogenous FBXO42 locus, and sequenced individual alleles using a TA cloning-like method.

### Xenograft tumors

All *in vivo* experiments were conducted in accordance with the NIH Guide for the Care and Use of Experimental Animals and with approval from the Fred Hutchinson Cancer Research Center, Institutional Animal Care and Use Committee (Protocol 1457). 100,000 GSCs were orthotopically xenografted into a single frontal cerebral hemisphere or in flanks of HSD:athymic nude Foxn1nu mice (#069, Envigo). For Dox-FBXO42 experiments, two days prior to tumor cell implant, 2mg/ml doxycycline with 5% sucrose (w/v) was added to the mouse drinking water.

## Data availability statement

The RNA-seq data that support the findings of this study are openly available in the NCBI Gene Expression Omnibus at https://www.ncbi.nlm.nih.gov/geo/, GEO accession number GSE213269 (token: cfqhwiemrfifrqf). Other data that support the findings of this study are available in the Supporting Materials of this article.

## Ethical statement

We followed the guidelines set by the Fred Hutchinson Cancer Center Institutional Review Office for De-identified Human Specimens and/or Data, which categorizes the studies presented here as Research Not Involving Human Subjects as detailed by the Institutional Review Board’s Human Subjects Research Determination Form.

